# Scaling and democratising structure-based protein function prediction with metagenomic-deepFRI

**DOI:** 10.64898/2026.04.27.720983

**Authors:** Valentyn Bezshapkin, Filip Schymik, Piotr Kucharski, Paweł Szczerbiak, Jakub W. Wojciechowski, Lukasz M. Szydlowski, Tomasz Kosciolek

## Abstract

High-quality protein structure models have become widely available, offering insights into protein function, yet they remain underutilized. Here, we introduce metagenomic-deepFRI, a framework incorporating structural templates into functional annotation pipelines at speeds comparable to sequence-alignment methods. Notably, structural features improved GO term prediction confidence and Information Content by up to 50%. Applied to metagenomic datasets, the framework achieves nearly 90% annotation coverage, enabling protein function inference without explicit orthology-based transfer.

## Main

Proteins underpin the functioning of all living matter, from maintaining cell structural integrity to catalysing metabolic reactions. While high-throughput metagenomic sequencing has revealed an immense diversity of proteins, driving an exponential expansion of public databases^1^, functional characterization lags significantly behind sequence acquisition. Swiss-Prot, a manually curated and annotated subset of UniProtKB, represents only around 0.3% of sequences in the database [https://www.uniprot.org/uniprotkb/statistics, release 2026_01 of UniProtKB, published on Wed Jan 28 2026.2]. Numerous computational approaches have been proposed to bridge this gap. For a long time, a significant part of such methods utilized only protein sequences. However, protein function depends primarily on its three-dimensional structure, which only recently became accessible for a large number of proteins due to the rise of deep learning models such as AlphaFold^2,3^ or ESM^4^. While such methods are a milestone for structural biology, their computational complexity hinders their use on metagenomic scale.

DeepFRI was developed to leverage protein structure for function prediction using graph convolutional networks^5^. Although its full structure-based mode may be underutilised for large-scale analyses due to the cost of predicting, storing and retrieving structures, it has nonetheless demonstrated substantial value in diverse biological contexts.

Applied to 1,070 infant gut metagenomes from the DIABIMMUNE cohort, deepFRI achieved 99% annotation coverage of 1.9 million assembled genes, compared to 12% by the orthology-based eggNOG^6–8^ — with 70% concordance between the two methods, establishing it as a viable complement to homology-based approaches^9^. Beyond metagenomics, structure-based annotations from deepFRI resolved functions in new and well-known environments, where sequence homology alone could not^10,11^. For example, in microbial fuel cell communities, it identified unique electron transport proteins in an electrogenic *Actinotalea* species^12^. In the representatives of the human gut methanoarchaea, deepFRI annotated previously uncharacterised genes, including a putative oxalyl-CoA decarboxylase and species-specific nucleotide-sugar transporters^13^. For newly-discovered bacterial species isolated from the International Space Station, the deepFRI approach helped identify adaptations to space conditions such as mechanosensitive channel proteins, S-layer oxidoreductases, and metallopeptidases with consistent glycine enrichment across all five species, none of which were detected by standard annotation pipelines^14^. A recent large-scale analysis integrating structures from the AlphaFold database^15^, ESMAtlas^4^, and the Microbiome Immunity Project^16^ demonstrated that high-level biological functions localise in specific regions of the protein structure space and that novel folds do not necessarily correspond to unknown functions^17^. Collectively, these studies demonstrate a clear demand for scalable, structure-aware annotation of metagenomic data. DeepFRI has proven capable of meeting this demand, but its structure-based mode remains underutilised at scale. Three barriers account for this. First, obtaining structures for millions of metagenomic proteins through de novo prediction (e.g. AlphaFold2) is computationally prohibitive. Second, even where predicted structures exist in public databases, storing and querying them efficiently at the scale of hundreds of millions of entries presents a significant data management challenge. Third, the original deepFRI implementation itself introduces inference overhead that compounds at the metagenomic scale. To address these barriers, we developed metagenomic-deepFRI (mdF), a computational pipeline that bypasses de novo structure prediction entirely by retrieving structures of homologous proteins from existing databases and transferring their structural information onto the query (Fig. 1A). Rather than predicting or storing structures for each query protein, the pipeline searches compressed structure databases using FoldComp^18^ and MMseqs2^19^, performs global sequence alignment with PyOpal^20,21^ to establish residue correspondence, and constructs a contact map that serves as input to the deepFRI graph convolutional network. For queries without a satisfactory structural match, the pipeline falls back to deepFRI’s sequence-only convolutional neural network. Inference is accelerated through ONNX (https://github.com/microsoft/onnxruntime), providing substantial speedup over the original implementation (Supplementary Fig. 1).

**Fig. 1.**
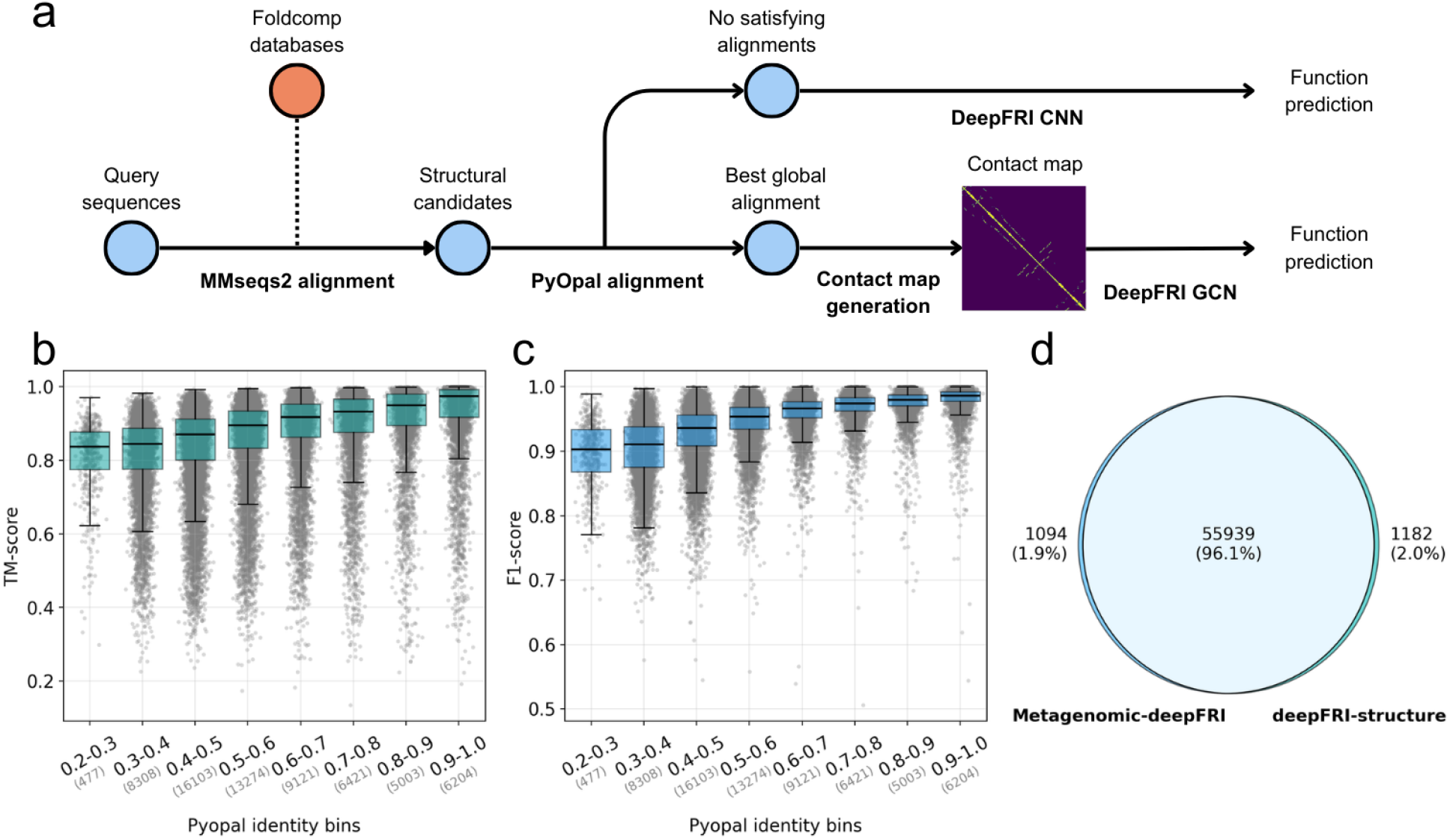
Pipeline overview and robustness of performance. **a**, Overview of metagenomic-deepFRI pipeline. **b**, Sequence identity vs TM-score between hit and reference proteins. **c**, Sequence identity vs F1 score of reconstructed contact map (gc parameter of 2). **d**, Number of molecular function GO terms overlapping between metagenomic-deepFRI and deepFRI using the reference structure with no identity handicap.

### Protein diversity benchmark

To initially validate the framework, we benchmarked metagenomic-deepFRI on a diverse set of proteins to ensure that it works robustly for different targets. We constructed a *Protein Diversity Dataset* (based on the Protein-structure-landscape^17^) that encompasses 50,000 non-redundant sequences with diverse folds. Those structures are high-quality predictions, originally from the AFDB50 database^10^ and are treated as a ground truth, to be compared with structures retrieved by the mdF algorithm.

We observe that our search retrieves structures similar to the ground truth, with a median TM-score exceeding 0.8 across the entire range of sequence identities (Fig. 1B), which is well above the 0.5 threshold typically indicative of a shared fold^22^. While DeepFRI remains robust when using lower-quality structural matches, high-quality structural priors significantly improve confidence scores and enable more granular GO terms annotations^23,24^ (Fig 1B, original deepFRI paper). The retrieved structure is then transformed into a contact map for the query protein. To handle insertions (gaps in alignment) and adapt the contact map as an input for the deepFRI, the pipeline imputes missing regions by enforcing local backbone connectivity (Supplementary Fig. 2). As shown in Figure 1C, the resulting contact maps achieve high accuracy relative to the ground truth, maintaining a median F1 score of ∼0.9 even in the ‘twilight zone’ of 20-30% sequence identity. This confirms metagenomic-deepFRI can effectively utilize structural information from remote homologs.

### Reference proteomes benchmark

To validate the framework on the function prediction task, we benchmarked metagenomic-deepFRI using five bacterial reference proteomes (Supplementary Table 1) from Uniprot with AlphaFold2-predicted structures as the ground truth^15^,^2^. We hypothesized that for those highly studied proteomes, mdF should be able to match many homologous structures and perform similar function predictions to standard structure-informed deepFRI, but without explicitly supplying the structures (high structural coverage demonstrated in Supplementary Fig. 3). Both methods showcased high prediction coverage (Supplementary Fig. 4). We observed a 96.1% overlap between Molecular Function GO terms^23,24^ predicted using our retrieval-based approach and those derived from dedicated structure predictions (Figure 1D), and overlaps of 94.7% and 89.7% when only accepting structural templates of maximum identity of 0.9 and 0.5, respectively (Supplementary Fig. 5). Figure 1D is built using no additional filtering of identity, other than removing self hits. With a median concordance index of 1.0 (Supplementary Fig. 6) per protein, metagenomic-deepFRI demonstrates that it can approximate the accuracy of a full structure-supplied run at a fraction of the computational cost. We also demonstrate that the structure retrieval of mdF significantly elevates the confidence scores, especially for the molecular function namespace of GO, reaching a median score above the typical high-confidence threshold (Supplementary Fig. 7).

### MAG benchmark

To test the utility of metagenomic-deepFRI for novel uncultivated bacterial genomes, we prepared a benchmark consisting of proteins from 50 high-quality metagenome-assembled genomes (MAGs)(Supplementary Table 2). To maximise structural coverage for these difficult targets, full AFDBv4 and clustered ESM databases were used (Supplementary Fig. 8), and minimum identity thresholds were reduced to 0.2 (Supplementary Fig. 9). First of all, incorporating structural information resulted in more specific functional annotations compared to a sequence-only model. We observed a significant increase in the Information Content (IC) of the predicted terms, particularly in Biological Process (median IC increase from 3.6 to 5.4) and Molecular Function (5.1 to 6.3) (Fig. 2A). Second, the median confidence score of deepFRI annotations was higher across all ontologies (BP, MF, and CC) compared to sequence-only prediction (Fig 2B). Even though the score increases were slightly less prominent compared to the reference proteomes benchmark (Supplementary Fig. 7), they still reached the median score, enabling high-confidence functional inference for many difficult targets.

**Fig. 2.**
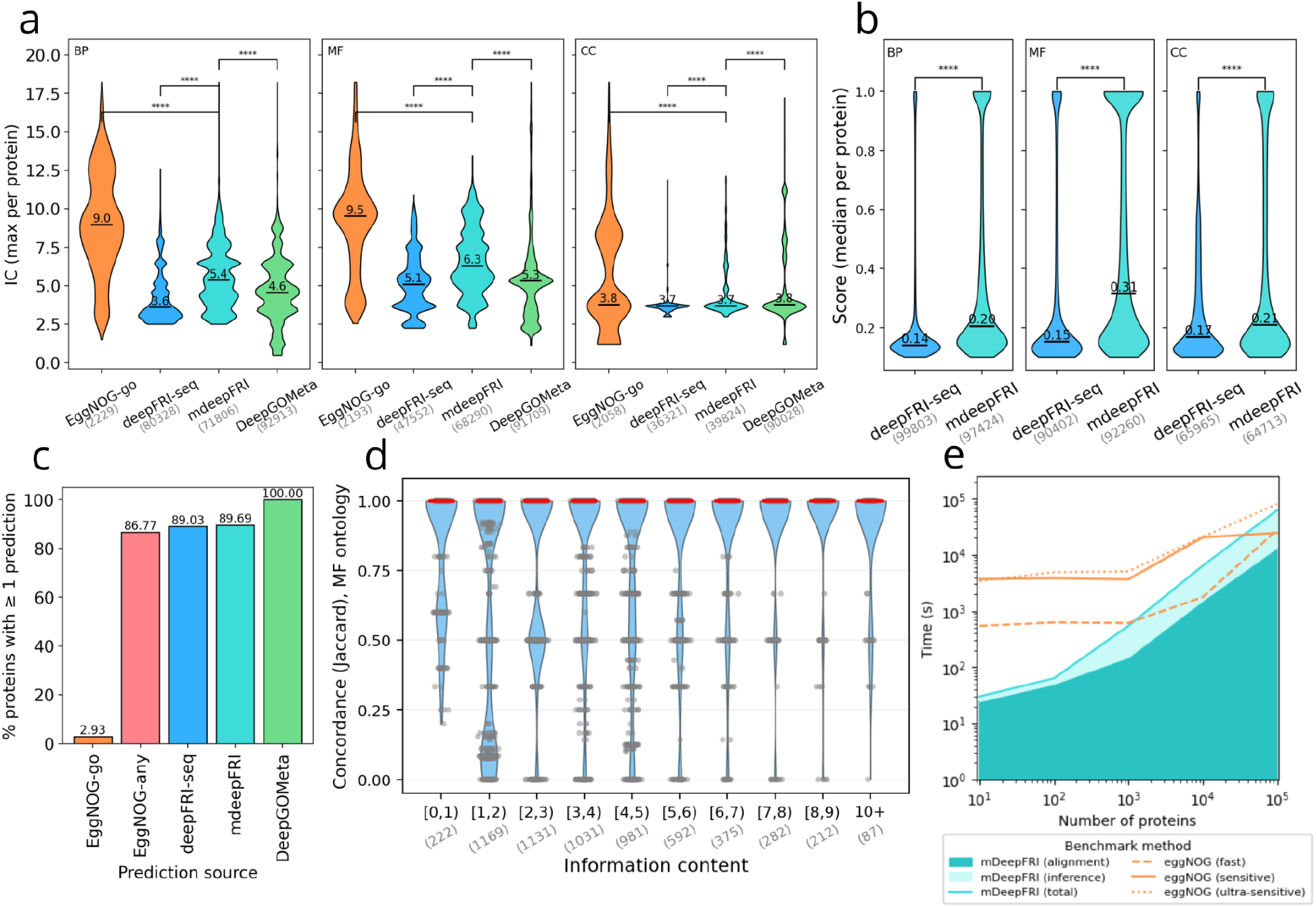
Function prediction performance comparisons. **a**, Per protein maximum IC values between methods (deepFRI-seq includes predictions of score 0.2 or more, and both metagenomic-deepFRI and DeepGOMeta of score 0.3 or more)(Mann-Whitney U-test, p < 1e-4). **b**, Score increase from metagenomic-deepFRI (Mann-Whitney U-test, p < 1e-4). **c**, Annotation coverage between methods (same score filtering as in point a). **d**, GO-term concordance between EggNOG-go and metagenomic-deepFRI for Molecular Function (other sub-ontologies shown in Supplementary Fig. 10 and 11). **e**, Method computational scaling comparison.

In comparison to the community standard EggNOG-mapper^6–8^, a distinct trade-off is observed. While EggNOG provides highly specific GO annotations (median IC > 9.0), it is limited by a severe bottleneck in coverage, annotating only 2.93% of the metagenomic proteins in our dataset with the standardized GO ontology. In contrast, metagenomic-deepFRI maintains a high coverage (89.69%), effectively bridging the gap between high-throughput sequencing and functional characterisation (Figure 2c). Only when additionally accepting annotations outside the GO ontology (Supplementary Note 1) EggNOG comes close to matching mdF in annotation coverage.

We also compared mdF against DeepGOMeta^25^, a recent GO prediction tool designed for metagenomic data that relies on protein sequence alone. To ensure a fair comparison, we applied the same GO propagation and coverage filtering to all three tools (see methods section on concordance analysis). DeepGOMeta annotates the entire MAG dataset (Fig. 2c), but the terms it predicts are less specific than those produced by mdF, as reflected by lower Information Content scores (Fig. 2a).

Thus, while EggNOG excels in precision for known homologs, DeepGOMeta in coverage, metagenomic-deepFRI is a first scalable solution (Fig 2e) offering integration of the protein structure information for function prediction, resulting in more specific annotations for a broader range of proteins.

### Conclusion

In summary, metagenomic-deepFRI enables scalable, structure-informed protein function prediction without *de novo* structure prediction. We demonstrate that retrieving and transferring structural information from homologous proteins approximates the accuracy of deepFRI supplied with dedicated structures. On metagenome-assembled genomes, the pipeline produces more specific annotations than a recent sequence-based method, while covering a larger fraction of proteins with GO terms than the orthology-based EggNOG-mapper. Several limitations should be noted. First, some protein sequences may lack homologs in public databases entirely, or the available structures may be of insufficient quality, both of which limit potential improvements to function prediction with mdF. As structural resources grow, annotation quality may improve correspondingly, but proteins from underrepresented taxonomies or with novel folds may lack suitable structural templates nevertheless. Second, contact maps remain an approximation of a three-dimensional structure, and with the added effects of alignment-based contact inference and the generated-contacts heuristic, they may induce noisy signals. Third, metagenomic-deepFRI inherits the limitations of the original model, such as a fixed vocabulary of 4,939 GO terms and the legacy architecture.

It is worth noting that the structure retrieval and contact map transfer components are model-agnostic, and any structure-based function prediction could leverage this part of the pipeline. In this sense, metagenomic-deepFRI is less a single tool than a general strategy for making structure-based annotation accessible at the metagenomic scale.

## Methods

### Computational pipeline

We developed a computational pipeline integrating structural database searching with the deepFRI framework for predicting protein function. The workflow consists of three distinct stages: structure retrieval and alignment; contact map construction; and deepFRI inference.

The pipeline begins by querying the input protein sequence against structural databases using MMSeqs2^19^. By default, PDB100 database is searched first, but other user-provided databases (e.g. AlphaFold Protein Structure Database, ESM Metagenomic Atlas) can be used (Supplementary Fig. 8). To enable efficient storage of large structural databases, we require all databases to be in the FoldComp^18,19^ format. After the search, we consider both alignment coverage and identity, and matches not meeting the default thresholds of 0.9 coverage and 0.5 identity are discarded. The top k hits (default: 5) from MMSeqs2 search results are then subjected to the global alignment using PyOpal (Python bindings for the Opal library implementing the Needleman-Wunsch algorithm^26^) and the best-scoring protein becomes the structural template, unless, after the global alignment, it does not meet the coverage and identity thresholds, then it is discarded.

Structural information in metagenomic-deepFRI is represented as a contact map (a binary distogram) that assumes a structural proximity of residues closer than 6Å. The alignment string from PyOpal is used to identify both matching and mismatching residues, for which contact information is directly transferred to the contact map. Global alignment from PyOpal is necessary to properly infer the residue-level correspondence used to populate the contact map; suboptimal, local alignment could misplace contacts, negatively affecting structure representation. For insertions in the query, we generate artificial contacts (Supplementary Fig. 2), preserving the backbone representation of the protein. The process is further described in the Contact map construction section.

The processed data is then fed into the deepFRI network, predicting Gene Ontology terms. If a structural template is available, the Graph Convolutional Network branch is activated on the contact map. Otherwise, the pipeline defaults to the Convolutional Neural Network using just the sequence.

The models are implemented in ONNX format, which gives 2-12 times the speedup over the original TensorFlow implementation, depending on sequence length and network type (Supplementary Fig. 1).

### Contact map construction

The structural input to deepFRI is a binary contact map - a square matrix, where each position (i, j) equals 1 if residues i and j are within a distance threshold. To construct a contact map, the pipeline proceeds in three steps: coordinate extraction, contact map computation based on the template and alignment-guided transfer to the query.

For each structure retrieved from a reference database, C-alpha coordinates are extracted using Biotite^27^, yielding a coordinate matrix of size depending on the number of template residues. Non-standard residues are mapped to their counterparts (e.g. selenomethionine to methionine). Pairwise squared Euclidean distances are computed between all C-alpha atoms, and binary contacts are assigned to each pair with a distance below 6 angstroms. The template contact map is then transferred to the query using the global alignment from string from PyOpal, which encodes matches, insertions and deletions. Walking that alignment string, the pipeline builds an index mapping template positions to query positions. Matched columns are treated as a direct correspondence, while deletions are unmapped.

Insertions are query residues with no structural information in the template. To prevent these from becoming isolated nodes in the graph convolutional network, the pipeline generates artificial contacts connecting inserted residues to their k nearest neighbours in the sequence in both directions (default k = 2). This enforces local backbone connectivity, based on the prior that residues close in sequence tend to be close in three-dimensions as well. The effect of the k parameter on the contact map is shown in Supplementary Fig. 2. The final output is an *N_query*

× *N_query* symmetric binary matrix serving as the structure-derived input for the deepFRI GCN.

### Protein Diversity Dataset

To reliably test the architecture of metagenomic-deepFRI we constructed a Protein Diversity Dataset, composed of non-redundant proteins with diverse structures. To construct this dataset, we used the Protein Structure Landscape^17^ (version 1.0) to randomly select 50,000 high-quality AFDB50^10,17^ proteins (pLDDT >90) out of all 1,505,141 proteins available in the database. We constructed reference contact maps using the obtained structures for later comparison, and extracted the sequence from these files for subsequent deepFRI runs.

### Contact map reconstruction benchmark

To test if metagenomic-deepFRI correctly reconstructs intermediate representations during its pipeline, we compared how similar are the structures and contact maps derived by our algorithm with the reference ones.

Metagenomic-deepFRI (version 1.1.8) was run on 50,000 sequences from the Protein Diversity Dataset, performing a MMseqs2 (version 13-45111+ds-2) search against the full AlphaFold v4 (https://foldcomp.steineggerlab.workers.dev/afdb_uniprot_v4 foldcomp version, uploaded 13, 14 Dec 2022). To investigate more distant, lower-identity matches and remove trivial pairings, self-hits were filtered out (same ID of query and match protein) and matches at various homology levels were picked using identity bins. Those bins were identity ranges formed between 0 and 1, with a 0.1 step. For each query protein, we selected a random hit within each bin. Those hits then followed a standard procedure of Pyopal (version 0.7.3) alignment, yielding structures and contact maps, to be compared with the reference. Precise parameters are described in the Supplementary Note 3.

The structures retrieved by metagenomic-deepFRI were saved and compared with the reference structures from the Protein Diversity Dataset using USAlign^28^ with the default settings. To compare the paired contact maps we used the F1-score, which is the harmonic mean of precision and recall. To differentiate between fragments of the contact map built from the structural information and artificial contacts, two derivative F1-scores were developed. Synthetic F1-score, which compares only artificial contacts, and inherited F1-score, which compares only the contacts inherited from the 3D structure.

### Reference proteomes dataset

To compare metagenomic-deepFRI’s performance with deepFRI supplied with generated structures, we chose five bacterial reference proteomes as a benchmarking set. The model organisms were selected for their high predicted structure coverage and phylogenetic diversity, as shown in the Supplementary Table 1.

A total of 14,895 modelled structures were obtained from the v6 version of the database (https://alphafold.ebi.ac.uk/download#proteomes-section, https://ftp.ebi.ac.uk/pub/databases/alphafold/latest/, uploaded 2025-08-24). We extracted sequences from these files using the Bio.PDB package (Biopython^29^ version 1.84). Those sequences were then used to run deepFRI^5,29^ (version 1.0.0) without any structural information, to enable comparison with structure-informed runs. To enable a fair comparison, we used a modified deepFRI implementation that uses ONNX models (https://github.com/microsoft/onnxruntime).

The downloaded structural models were used to run deepFRI in the same way as the sequence-only deepFRI discussed previously, but this time with structural information. Lastly, metagenomic-deepFRI (version 1.1.8) was used on the proteomes. To ensure structural coverage we searched the default PDB (https://wwwuser.gwdg.de/∼compbiol/colabfold/pdb100_230517, last accessed 09.03.2026) database as well as the full AFDB (https://foldcomp.steineggerlab.workers.dev/afdb_uniprot_v4, uploaded 13, 14 Dec 2022) and ESM Atlas clusters (https://foldcomp.steineggerlab.workers.dev/highquality_clust30, uploaded 13 Mar 2025) FoldComp databases. Similarly to the contact map reconstruction benchmark, retrieving the exact structure of the query protein was avoided by excluding self-hits (same ID of query and match protein). Since more distant matches are beneficial to examine, we kept some lower identity hits, by independently removing hits above 0.5 identity, above 0.9 identity and with no additional filtering, forming in effect 3 subsets.

Those MMSeqs2^19^ (version 13-45111+ds-2) matches followed the rest of the standard pipeline, being used for contact map generation and function prediction. Precise parameters are described in the Supplementary Note 3.

### Concordance analysis

Concordance^9^ is a metric that measures the degree of agreement between two sets of go-terms, independent of semantic similarity. The metric is calculated using the Jaccard index. The number of GO terms present in both sets is divided by the total number of unique GO terms. For perfectly overlapping sets, the value of concordance is equal to 1. The lower the overlap, the closer to 0 is the metric.

To make the comparison fair, we first filter out GO terms^23,24^ that deepFRI cannot predict and then propagate the GO terms in all the sets under consideration. Propagation takes all remaining predictions for a given protein and, based on the Gene Ontology graph in the .obo (http://geneontology.org/docs/download-ontology/go-basic.obo, release 2025-10-10) file format, adds all the parent terms down to the root using the “is_a” relationship. If present, the scores are inherited based on the strongest child evidence. If a parent term has many child terms with different scores, the strongest evidence is the highest score among them. The concordance is then stratified using GO aspects and Information Content bins.

### MAG dataset

To test metagenomic-deepFRI in a realistic scenario, where the user might use novel microbial sequences without known structures as input, we prepared a set of metagenome-assembled genomes (MAGs) from the human gut microbiome as a challenging function prediction task. MAGs are draft microbial genomes reconstructed from sequencing data by joining reads into contigs and then binning those into genome representations.

Fifty metagenomic assembly genomes were selected from the Unified Human Gastrointestinal Genome^30^ (UHGG) (genomes-all_metadata.tsv, last modified 28 September 2023, 13:23, EBI FTP). First, we filtered the data to include only high-quality entries by enforcing a completeness threshold of more than 95%, a contamination threshold of less than 1%, and a number of contigs threshold of less than 100. As only the species representatives have EggNOG and sequence information available, we selected those species representatives that are MAGs. The selected genomes are shown in the Supplementary Table 2. Sequence information and EggNOG annotation files were downloaded from the EBI ftp (‘https://ftp.ebi.ac.uk/pub/databases/metagenomics/mgnify_genomes/human-gut/v2.0.2/species_catalogue/{id}/{id}/genome/{id}.faa/{id}_eggNOG.tsv‘, last modified 2021-12-07).

These sequences were then used to run deepFRI using ONNX models for fair comparison. To gather as much structural information as possible, metagenomic deepFRI was run against the PDB (https://wwwuser.gwdg.de/∼compbiol/colabfold/pdb100_230517, last accessed 09.03.2026), afdb_uniprot (https://foldcomp.steineggerlab.workers.dev/afdb_uniprot_v4, uploaded 13, 14 Dec 2022) and ESM Atlas clusters (https://foldcomp.steineggerlab.workers.dev/highquality_clust30, uploaded 13 Mar 2025). FoldComp databases and hits of any identity were accepted. The Pyopal (version 0.7.3) filter was liberal to maximise structural coverage for those difficult targets - minimum contact map identity was 0.2 and the minimum coverage was 0.9.

Gene Ontology term prediction was carried out with the standard procedure. Precise parameters are described in the Supplementary Note 3.

To compare the results with those of an alternative GO-term prediction tool, we ran DeepGOMeta^25^ (https://github.com/bio-ontology-research-group/deepgometa, commit fbbe33d) to analyse the additional structural information gained by metagenomic deepFRI (version 1.1.8) compared to a very recent sequence-only prediction tool. Since the default score-filtering threshold on the DeepGOWeb page is 0.3, we used this value in our analysis (https://deepgo.cbrc.kaust.edu.sa/deepgo/).

### Information Content

Our Information Content definition follows the original deepFRI paper^5^.

## Data availability

All source and intermediate files, along with scripts and plots are available via Zenodo at at 10.5281/zenodo.19738140.

## Code availability

Metagenomic-deepFRI is available under the BSD 3-Clause license via GitHub at https://github.com/Tomasz-Lab/metagenomic-deepFRI. Documentation is available at https://metagenomic-deepfri.readthedocs.io. The analysis notebooks are available via GitHub at https://github.com/Tomasz-Lab/metagenomic-deepfri-project.

## Ethics declarations

### Competing interests

None declared.

## Author contributions

V.B., F.S., T.K., J.W.W., P.S. designed the experiments; V.B., F.S., P.K. designed and implemented the algorithms and performed the benchmarks; F.S., V.B., J.W.W., P.S., T.K. analysed the results; J.W.W., L.M.S. prepared benchmark data; L.M.S. prepared biological interpretation of the data; P.K. made the original contact map alignment implementation and release of metagenomic-deepFRI; P.S. helped with algorithm design and test during the first release; F.S., V.B., J.W.W., T.K., P.S. wrote the paper; T.K. conceived and supervised the project; All authors have read and approved the final version of the paper.

## Acknowledgements

This research was in part conducted in the laboratory of Prof. Dr. Shinichi Sunagawa at Institute of Microbiology, ETH Zurich. The authors thank Prof. Sunagawa for the opportunity to carry out this work and for providing a supportive environment for independent research. The authors would also like to thank Prof. Aleksander Byrski, Dean of the Faculty of Computer Science at AGH University of Krakow, for his support, PhD supervision (F.S.), and fostering the bioinformatics community at the University that contributed to the development of this project. We would also like to thank Rund Tawfiq (Sano) and Kinga Zielińska (Małopolska Centre of Biotechnology, Jagiellonian University in Krakow) for their invaluable advice and feedback on this project.

T.K., F.S., J.W.W. are supported by the National Science Centre, Poland grant 2023/05/Y/NZ2/00080. This research was funded by the project of the Minister of Science and Higher Education “Support for the activity of Centers of Excellence established in Poland under Horizon 2020” on the basis of the contract number MEiN/2023/DIR/3796. We gratefully acknowledge Polish high-performance computing infrastructure PLGrid (HPC Center: ACK Cyfronet AGH) for providing computer facilities and support within computational grant no. PLG/2025/018263.

## Notes

https://github.com/Tomasz-Lab/Metagenomic-DeepFRI

https://metagenomic-deepfri.readthedocs.io/

https://doi.org/10.5281/zenodo.19738140

### Competing Interest Statement

The authors have declared no competing interest.

