## Supplementary Information for "Scaling and democratising structure-based protein function prediction with metagenomic-deepFRI"

**Supplementary Table 1:** Five reference proteomes used in the benchmark.

| Reference Proteome | Species | Phylum | Class | No. of Predicted Structures |
|---|---|---|---|---|
| UP000000625 | Escherichia coli | Pseudomonadota | Gammaproteobacteria | 4,370 |
| UP000000429 | Helicobacter pylori | Campylobacterota | Epsilonproteobacteria | 1,540 |
| UP000001584 | Mycobacterium tuberculosis | Actinomycetota | Actinomycetes | 3,991 |
| UP000008816 | Staphylococcus aureus | Bacillota | Bacilli | 2,888 |
| UP000000535 | Neisseria gonorrhoeae | Pseudomonadota | Betaproteobacteria | 2,106 |

**Supplementary Table 2:** MAGs used in the benchmark. Fasta files and EggNOG tsv files were downloaded from the EBI FTP. Contamination is on the scale from 0 to 100, we kept only genomes with less than 1% contamination.

| Species_rep | Completeness | Contamination | No. of contigs | Phylum | Class | Species |
|---|---|---|---|---|---|---|
| MGYG000000354 | 96.48 | 0.57 | 58 | Bacteroidota | Bacteroidia | CAG-485 sp001701295 |
| MGYG000000378 | 97.26 | 0.95 | 91 | Firmicutes_A | Clostridia | |
| MGYG000000412 | 99.38 | 0.29 | 30 | Proteobacteria | Gammaproteobacteria | Succinivibrio sp000431835 |
| MGYG000000513 | 98.42 | 0.35 | 53 | Campylobacterota | Campylobacteria | Helicobacter_B fennelliae |
| MGYG000000525 | 99.38 | 0.62 | 78 | Proteobacteria | Gammaproteobacteria | Duodenibacillus sp900767875 |
| MGYG000000530 | 97.18 | 0.72 | 73 | Firmicutes_A | Clostridia_A | PeH17 sp000435055 |
| MGYG000000534 | 96.86 | 0.05 | 85 | Bacteroidota | Bacteroidia | Prevotella sp900543975 |
| MGYG000000896 | 100 | 0.81 | 38 | Actinobacteriota | Coriobacteriia | Collinsella sp900540895 |
| MGYG000001130 | 95.16 | 0.81 | 59 | Actinobacteriota | Coriobacteriia | Collinsella sp900757615 |
| MGYG000001136 | 96.31 | 0.67 | 24 | Firmicutes_A | Clostridia | CAG-103 sp900543625 |
| MGYG000001178 | 98.1 | 0.69 | 84 | Firmicutes_A | Clostridia | Anaerosacchariphilus sp900553635 |
| MGYG000001605 | 100 | 0 | 41 | Actinobacteriota | Coriobacteriia | Enorma sp900538305 |
| MGYG000001664 | 96.92 | 0.75 | 96 | Bacteroidota | Bacteroidia | CAG-485 sp900760735 |
| MGYG000001745 | 99.33 | 0 | 26 | Firmicutes_A | Clostridia | Coprococcus sp900548215 |
| MGYG000001785 | 97.33 | 0 | 54 | Firmicutes | Bacilli | UMGS1109 sp900548675 |

| | | | | | | |
|---|---|---|---|---|---|---|
| MGYG000001810 | 95.52 | 0.61 | 56 | Firmicutes_A | Clostridia | Mediterraneibacter sp002314255 |
| MGYG000001927 | 99.15 | 0 | 74 | Bacteroidota | Bacteroidia | Prevotella sp900547005 |
| MGYG000001987 | 97.99 | 0 | 55 | Firmicutes_A | Clostridia | UMGS363 sp900543105 |
| MGYG000001993 | 100 | 0.81 | 90 | Actinobacteriota | Coriobacteriia | Collinsella sp900754275 |
| MGYG000002019 | 97.32 | 0 | 23 | Firmicutes_A | Clostridia | Eubacterium_R sp000431535 |
| MGYG000002087 | 100 | 0.81 | 8 | Actinobacteriota | Coriobacteriia | Collinsella sp900545445 |
| MGYG000002260 | 97.65 | 0 | 66 | Firmicutes_A | Clostridia | Eubacterium_R sp900540235 |
| MGYG000002591 | 98.02 | 0.62 | 28 | Proteobacteria | Gammaproteobacteria | CAG-521 sp902388655 |
| MGYG000002645 | 98.2 | 0 | 23 | Firmicutes_C | Negativicutes | Veillonella dispar_A |
| MGYG000002852 | 96.77 | 0.81 | 89 | Actinobacteriota | Coriobacteriia | Collinsella sp900543515 |
| MGYG000002911 | 100 | 0.81 | 44 | Actinobacteriota | Coriobacteriia | Collinsella sp900541045 |
| MGYG000002932 | 100 | 0.81 | 47 | Actinobacteriota | Coriobacteriia | Collinsella sp900541055 |
| MGYG000003007 | 100 | 0.81 | 62 | Actinobacteriota | Coriobacteriia | Collinsella sp900541025 |
| MGYG000003012 | 97.72 | 0.32 | 48 | Firmicutes_A | Clostridia | Blautia_A sp900540785 |
| MGYG000003122 | 100 | 0.81 | 34 | Actinobacteriota | Coriobacteriia | Collinsella sp900541235 |
| MGYG000003129 | 98.23 | 0.91 | 39 | Firmicutes_A | Clostridia | Anaerococcus octavius |
| MGYG000003329 | 97.11 | 0.64 | 53 | Firmicutes_C | Negativicutes | Veillonella sp900550455 |
| MGYG000003361 | 98.79 | 0 | 16 | Actinobacteriota | Coriobacteriia | Collinsella sp900556495 |
| MGYG000003376 | 98.88 | 0.16 | 19 | Fusobacteriota | Fusobacteriia | Sneathia sanguinegens |
| MGYG000003466 | 97.2 | 0 | 66 | Spirochaetota | Spirochaetia | Treponema_D sp900767955 |
| MGYG000003583 | 98.66 | 0 | 98 | Firmicutes_A | Clostridia | Ruminococcus_C sp900770195 |
| MGYG000003929 | 99.33 | 0 | 18 | Firmicutes_A | Clostridia | CAG-590 sp900544905 |
| MGYG000004021 | 97.82 | 0.01 | 37 | Bacteroidota | Bacteroidia | UBA11471 sp900542765 |
| MGYG000004099 | 97.32 | 0 | 13 | Firmicutes_A | Clostridia | Eubacterium_R sp000433975 |
| MGYG000004133 | 98.66 | 0 | 94 | Firmicutes_A | Clostridia | |
| MGYG000004181 | 98.99 | 0.81 | 86 | Actinobacteriota | Coriobacteriia | Collinsella sp002232035 |
| MGYG000004231 | 95.41 | 0.67 | 93 | Firmicutes_A | Clostridia | |
| MGYG000004321 | 96.87 | 0 | 34 | Firmicutes | Bacilli | Streptococcus constellatus |
| MGYG000004402 | 96.43 | 0 | 67 | Bacteroidota | Bacteroidia | RC9 sp000432515 |
| MGYG000004451 | 100 | 0.11 | 25 | Firmicutes | Bacilli | CAG-313 sp003539625 |
| MGYG000004625 | 100 | 0.81 | 29 | Actinobacteriota | Coriobacteriia | |

| | | | | | | |
|---|---|---|---|---|---|---|
| MGYG000004715 | 99.29 | 0.71 | 80 | Firmicutes_A | Clostridia | Eubacterium_T saphenum_A |
| MGYG000004847 | 97.99 | 0.89 | 50 | Firmicutes_A | Clostridia | UMGS1670 sp902406135 |
| MGYG000004866 | 96.66 | 0.68 | 80 | Firmicutes_A | Clostridia | Gemmiger variabilis_B |
| MGYG000004880 | 97.04 | 0.19 | 39 | Bacteroidota | Bacteroidia | |

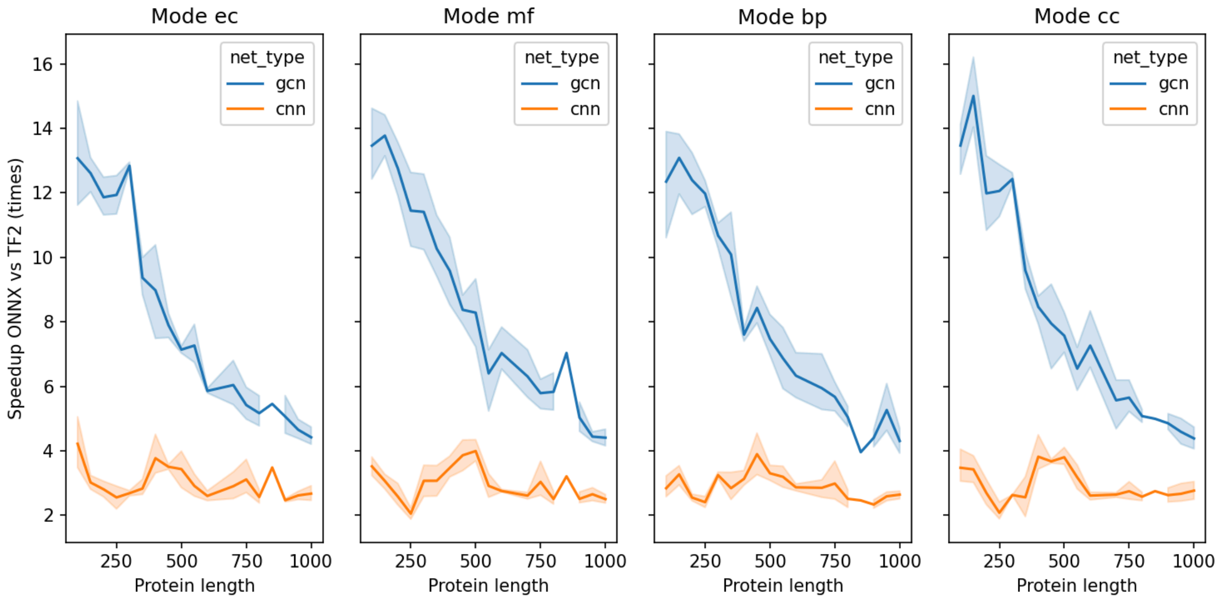


**Supplementary Figure 1:** ONNX model speedup. The original DeepFRI models were trained and distributed as TensorFlow 2 / Keras models (.hdf5). To reduce the dependency footprint and improve inference speed, all eight models (CNN and GCN variants for EC, MF, BP, and CC) were converted to ONNX format using the tf2onnx library. To validate the conversion, 100 randomly generated protein sequences (lengths 60-1000 aa) were passed through both the original TensorFlow and the converted ONNX models. For GCN models, random binary contact maps of matching dimensions were generated alongside the sequences. Inference wall-clock times were recorded for each protein on a single CPU core. The ONNX Runtime was consistently faster than TensorFlow 2 across all protein lengths and prediction modes. The speedup was most notable for the CNN models. GCN models also showed improvements, though the relative gains were smaller for longer sequences.

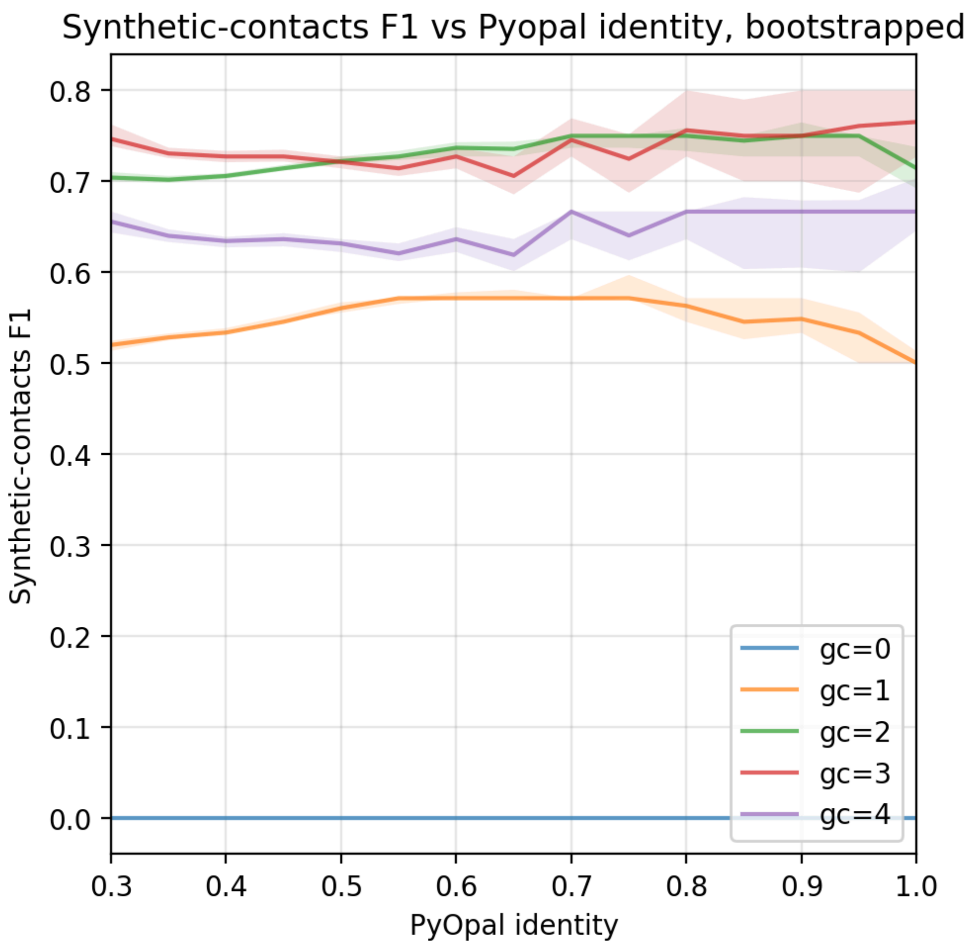


**Supplementary Figure 2:** This benchmark was performed on the Protein Diversity dataset. Contact maps were reconstructed from the same structures, changing only the --generate_contacts parameter (gc) from 0 to 4. Only residues for which structural information cannot be inferred were chosen for F1 calculation to explicitly focus on the effects of generated contacts. As a sanity check gc=0 is included, expectably showing the lowest possible scores since no missing information was filled. Both gc 2 and 3 perform very well, showing performance between 0.7 and 0.8 F1. gc=2 is set as a default value in mDeepFRI, since it performs more robustly than gc=3 (based on bootstrap confidence intervals).

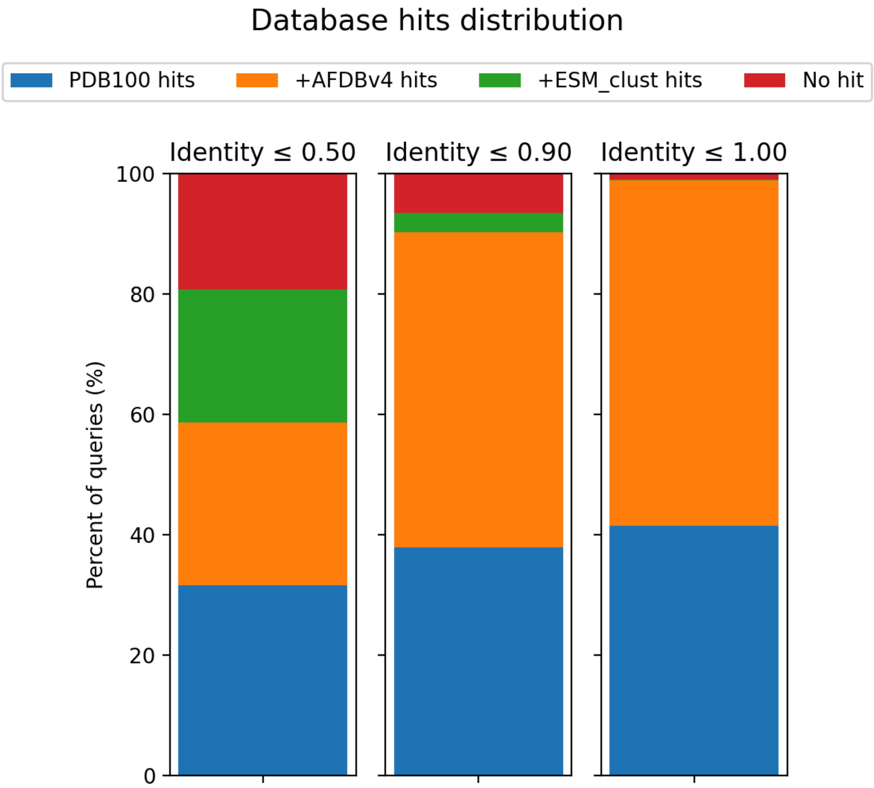


**Supplementary Figure 3:** Structural coverage in the Reference Proteomes benchmark. When all-identity hits were accepted, nearly all queries had a structure retrieved. As the pipeline was forced to only find more distant hits to the query, the coverage decreased.

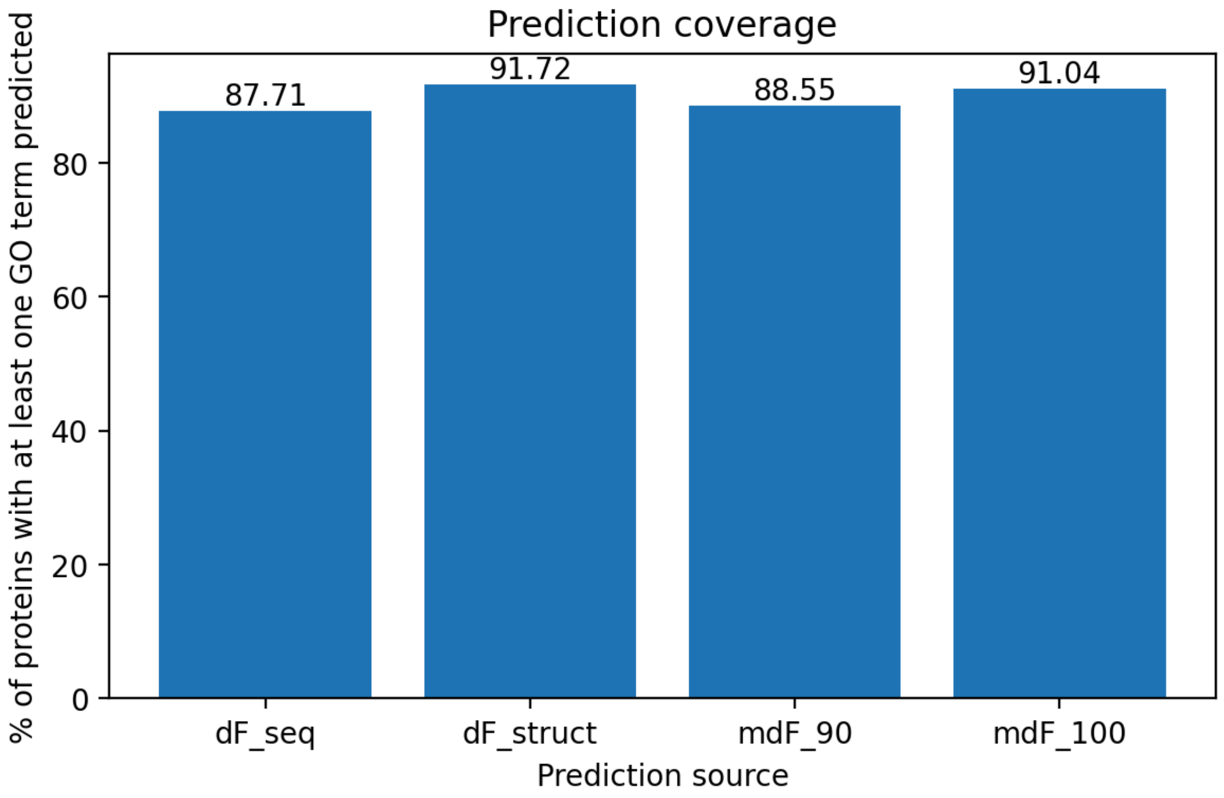


**Supplementary Figure 4:** This plot is based on the reference proteomes data. Coverage for 4 function prediction approaches. df_seq is deepFRI run on sequence only and filtered for score >= 0.2 dF struct is deepFRI with ground truth structures supplied and filtered for score >= 0.3. mdF_90 and mdF_100 are metagenomic-deepFRI runs, limited to a maximum sequence identity of 0.9 and no limit accordingly, filtered for score >= 0.3. MMSeqs2 minimum identity threshold was disabled. PyOpal contact map identity was set to 0.3. Same ID hits were discarded (query sequences find their own ID in the reference database). Top K filtering in mDeepFRI was disabled. The total number of proteins in the Reference Proteomes benchmark was 14895.

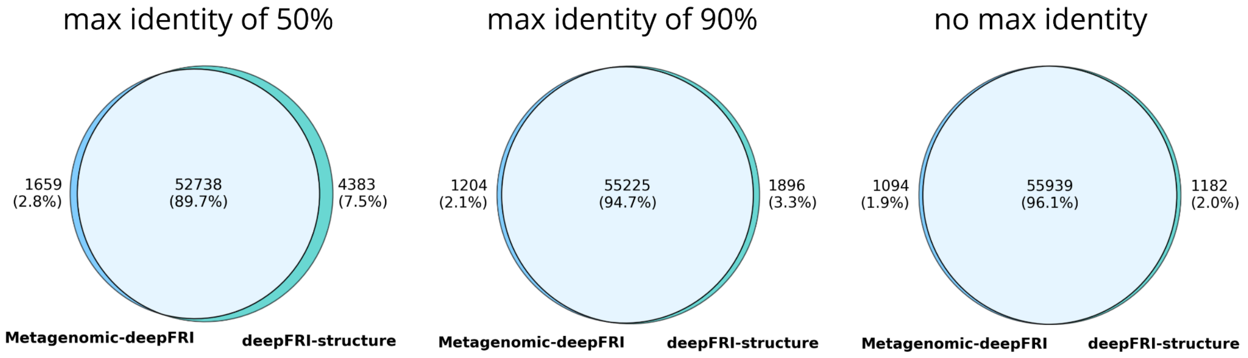


**Supplementary Figure 5:** Overlap change for different max identity thresholds when comparing MF GO terms predicted in the reference proteomes benchmark.

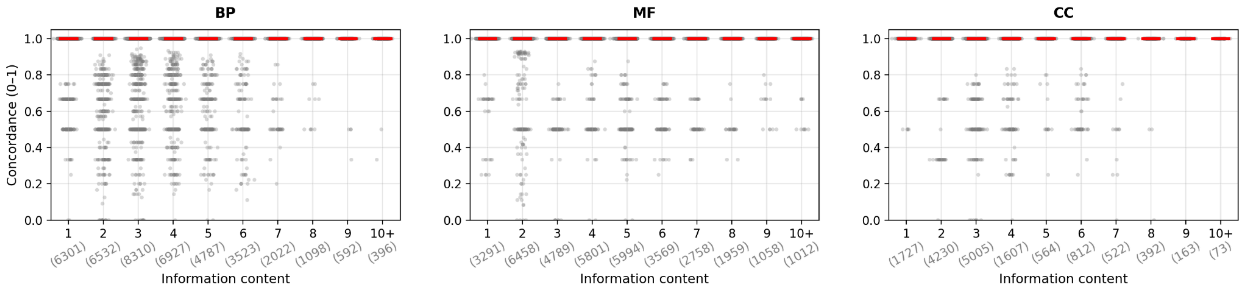


**Supplementary Figure 6:** Concordance of metagenomic-deepFRI and structure-supplied deepFRI on 5 reference proteomes. Both methods were filtered on a 0.3 score threshold and propagated before the comparison. Obsolete GO terms were discarded. The plot represents a boxplot with medians highlighted in red, boxes and whiskers concentrated around 1 and outliers visible on the graph.

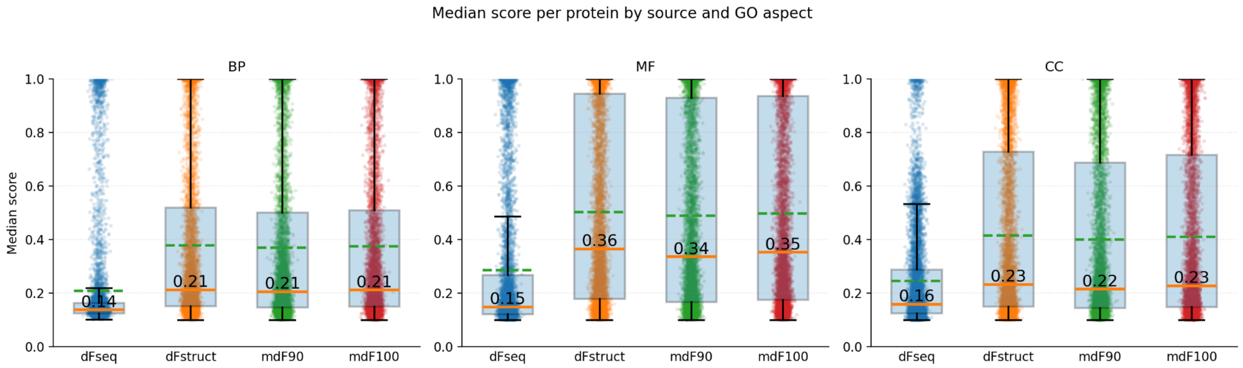


**Supplementary Figure 7:** Distributions of median scores per protein in the Reference Proteomes benchmark. The green dashed line is the mean, and the orange line is the median. No prior score filtering is applied. For structure-based predictions, using a GCN score of 0.3 and more is considered high confidence, and for sequence-only predictions, using a CNN score of 0.2 and more is considered high confidence. mdF_90 and mdF_100 are metagenomic-deepFRI runs, limited to a maximum sequence identity of 0.9 and no limit accordingly.

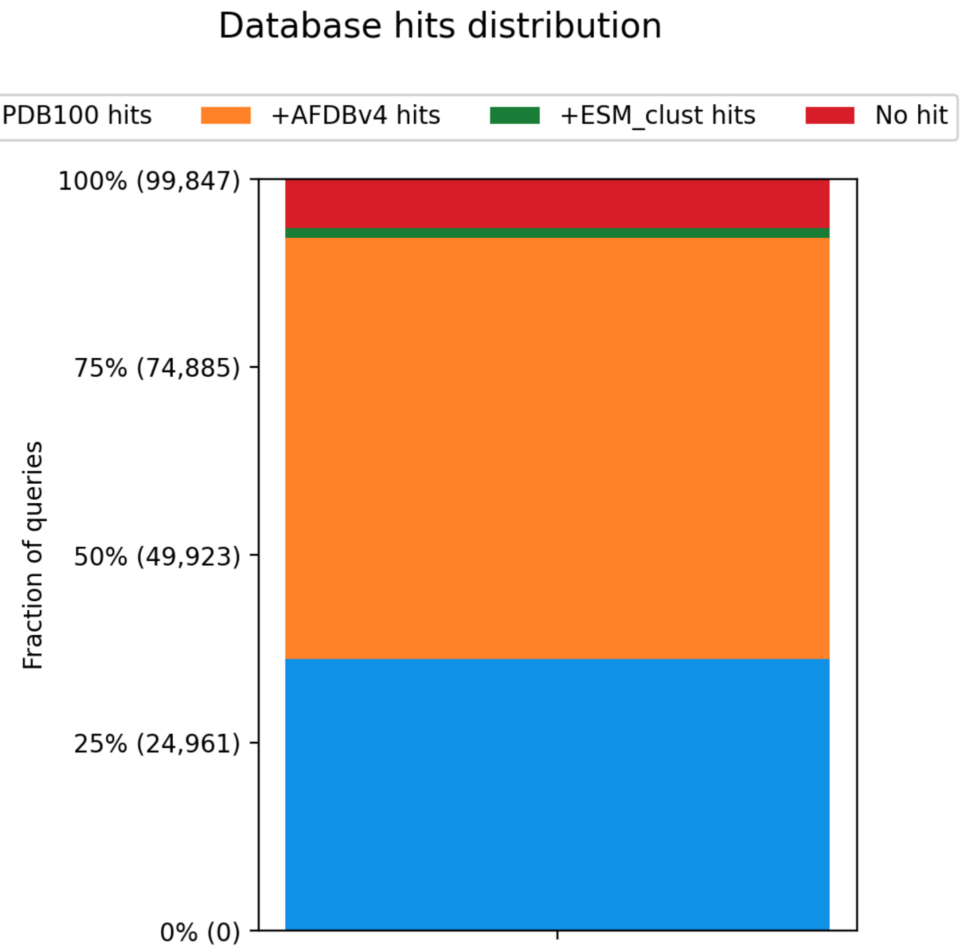


**Supplementary Figure 8:** This plot was built using the data from the MAG benchmark. Even for those more challenging metagenomic samples, high structural coverage can be reached by incorporating additional databases. Full AFDBv4 is a large reference database, but once set-up it can be reused for many experiments leveraging mDeepFRI.

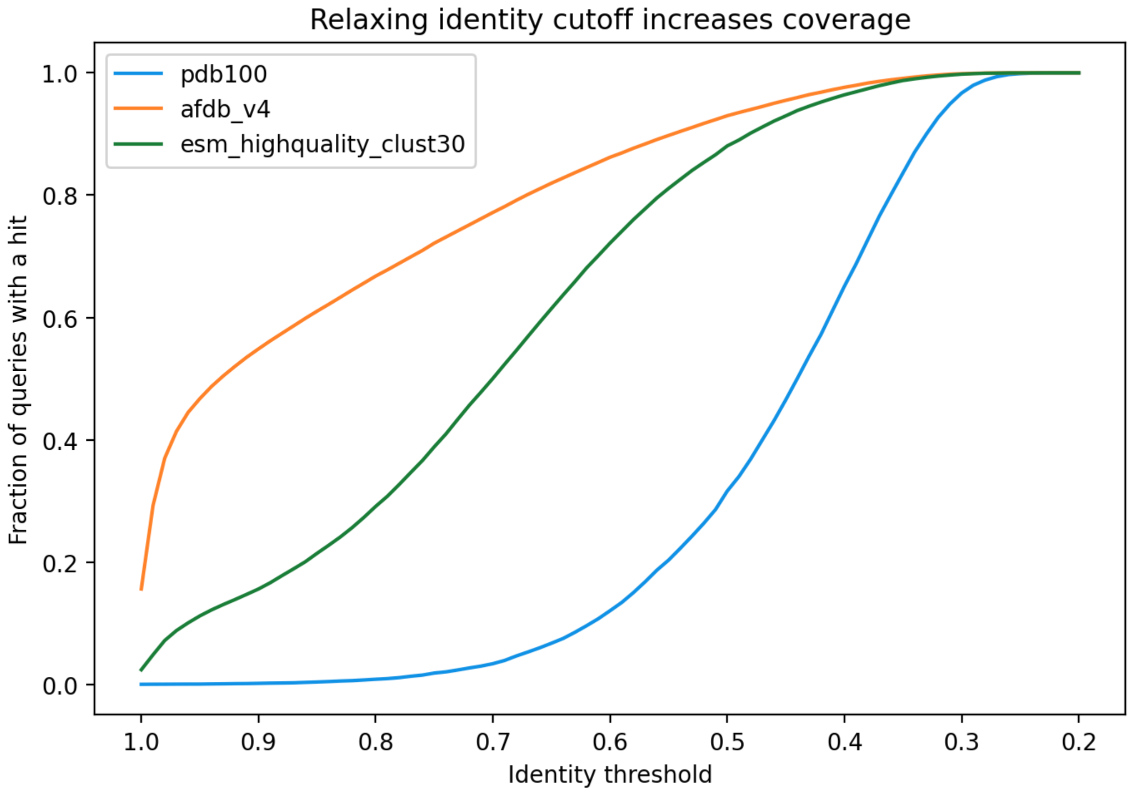


**Supplementary Figure 9:** This benchmark was performed on the MAG dataset. 100% on the y-axis are all hits for a given database for this mDeepFRI run. A sudden jump in hits occurs at a lower identity threshold for pdb, showing that for this database additional hits can be found below 0.5 identity (default). For the other two databases there is not a big benefit in going below 0.5 identity in this run. Users seeking high coverage for difficult targets, might want to lower their identity (and coverage) thresholds for significant increases in structural coverage.

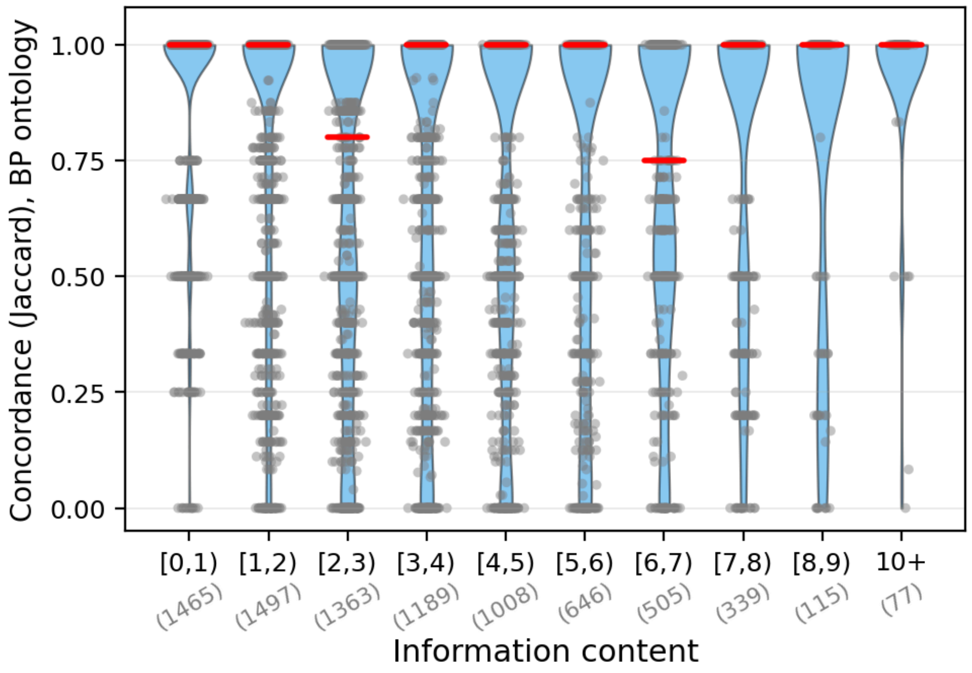


**Supplementary Figure 10:** GO-term concordance between EggNOG-go and metagenomic-deepFRI for Biological Process.

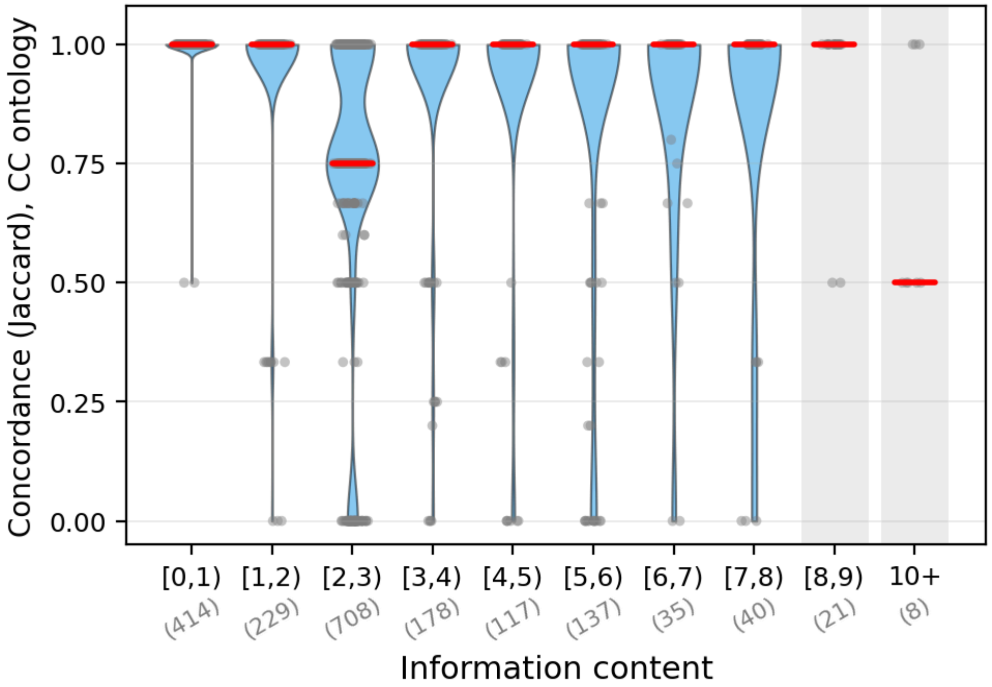


**Supplementary Figure 11:** GO-term concordance between EggNOG-go and metagenomic-deepFRI for Cellular Component.

### Supplementary Note 1

**EggNOG filtering**

Fig. 2c has a category for EggNOG-any annotations. Those are built from entries that among these columns have any entry: `COG_category Description Preferred_name GOs EC KEGG_ko KEGG_Pathway KEGG_Module KEGG_Reaction KEGG_rclass BRITE KEGG_TC CAZy BiGG_Reaction PFAMs`. Unless the only source of information is `COG_category` being “S”, `PFAMs` being `"^(?:DUF\d*|unknown|uncharacteri[sz]ed|hypothetical)$"` and `Description` being: `r"(?:^|\b)(protein of unknown function|hypothetical protein|uncharacteri[sz]ed protein|", r"domain of unknown function|unknown function)(?:\b|$)"`.

### Supplementary Note 2

**Metagenomic-deepFRI modified version**

We used a modified version of the public-facing metagenomic-deepFRI that extends the standard pipeline with several small changes to support systematic benchmarking and finer control over structure retrieval:

1. The modified version introduces an `--identity-bin` parameter that accepts multiple identity ranges (e.g., `0.80,0.90`). This allows systematic evaluation of how restricting MMseqs2 hit identity to specific ranges affects functional prediction quality. The original version uses a single identity threshold (`--mmseqs-min-identity`, default 0.5) with no upper-bound filtering.
2. The modified version accepts multiple `--generate-contacts` values in a single run, iterating over all combinations of (`identity-bin × generate-contacts`). The original version only supports a single `generate-contacts` value per run.
3. A new `filter_mmseqs_best_matches` function was added that operates on saved MMseqs2 TSV files. This enables: (a) filtering hits into identity bins (lower inclusive, upper exclusive), (b) removal of self-hits, (c) per-query hit selection via either top-bitscore or random sampling.
4. In the modified version, setting `--top-k 0` retains all MMseqs2 hits, rather than selecting only the `top-k` per query. The original version defaults to `top-k = 3`.
5. The added `--skip-prediction` flag allows running only the search, alignment, and contact map generation steps without GO-term prediction, which is useful for benchmarking contact map quality independently.
6. The added `--save-raw-alignments` flag exports gapped query and target sequences from PyOpal alignments in FASTA format, along with alignment scores in TSV format.

### Supplementary Note 3

**Metagenomic-deepFRI run settings**

All three runs (corresponding to three benchmarks used in the paper) used MMseqs2 for initial homology search with default sensitivity (5.7) and default maximum E-value (0.001). PyOpal was used for subsequent pairwise sequence alignment with gap-open penalty 10 and gap-extend penalty 1 (defaults). The key differences between runs are in the databases searched, the identity filtering strategy, and the contact map generation parameters.

***Run 1: GC Benchmark***
Purpose: systematic benchmarking of contact map generation across identity bins and generate-contacts values, without running GO-term prediction.

Input: 50K protein sequences from protein diversity landscape
Databases: AlphaFold Database UniProt v4 (afdb_uniprot_v4)
Skip PDB: Yes | PDB100 was not searched
MMseqs2 parameters:

- Sensitivity 5.7 (default)
- Max E-value 0.001 (default)
- Min identity 0.0 | No identity filtering at the MMseqs2 stage
- Min coverage 0.9 (default)
- Top-K 0 (all hits) | All passing MMseqs2 hits were retained |

Post-filtering:

- Identity bins: 10 bins: [0.00, 0.10), [0.10, 0.20), ..., [0.90, 1.00) | Each bin contains hits within a 10% identity range |
- Per-query selection | random (seed = 42) | One hit per query selected randomly within each bin |
- Drop self-hits Yes | Self-hits removed |

PyOpal alignment filtering:

- cmap-identity 0.0 | No identity filtering after PyOpal alignment
- cmap-coverage 0.0 | No coverage filtering after PyOpal alignment

Contact map:

- Generate contacts: 0, 1, 2, 3, 4 | All five gap-fill values tested |
- Angstrom threshold | 6 Å (default)

Prediction: Processing modes: None (skipped) | GO-term prediction was disabled
Output:

- Save contact maps | Yes
- Save structures | Yes | |
- Save raw alignments | Yes | Gapped alignments exported

This run produced 10 (`identity bins`) × 5 (`generate-contacts` values) = 50 output combinations, enabling evaluation of how homolog identity and contact map gap-filling interact.

***Run 2: Reference Proteomes***
Purpose: functional annotation of reference proteomes using a three-database hierarchical search with cumulative identity thresholds.

Input: 5 Reference proteome sequences
Databases: PDB100 → AFDB UniProt v4 → ESM highquality_clust30 | Three-database hierarchical search

Skip PDB: No | PDB100 was searched first
MMseqs2 parameters:

- Sensitivity | 5.7 (default)
- Max E-value | 0.001 (default)
- Min identity: 0.0 | No identity filtering at MMseqs2
- Min coverage | 0.9 (default)
- Top-K | 0 (all hits) | All passing hits retained

Post-filtering:

- Identity bins: [0.00, 0.50), [0.00, 0.90), [0.00, 1.01) | Cumulative bins with increasing upper bounds
- Per-query selection: topbits (default) | Best hit per query by bitscore
- Drop self-hits: Yes

PyOpal alignment filtering:

- Cmap-identity: 0.3 | Alignments with < 30% identity discarded
- Cmap-coverage: 0.9 | Alignments with < 90% query coverage discarded

Contact map:

- Generate contacts: 2

Prediction:

- Processing modes: BP, CC, MF | Biological Process, Cellular Component, Molecular Function

The three cumulative identity bins ([0.00, 0.50), [0.00, 0.90), [0.00, 1.01)) enable comparison of predictions when restricting to distant homologs only (< 50% identity), moderate homologs (< 90%), or including near-identical matches (< 101%, effectively all hits). Note that the upper bound of identity bins is exclusive, so 1.01 is used to include perfect 1.0 identity matches. Post-PyOpal filtering with 30% identity and 90% coverage ensures that only structurally meaningful alignments are used for contact map transfer. Only the [0.00, 1.01) bin accepting all hits is considered in the main text, the [0.00, 0.90) is referenced in Supplementary Figure 7 under the name mdF_90, and all three are compared in Supplementary Figure 5.

***Run 3: MAGs***

Purpose: functional annotation of metagenomic assembled genomes (MAGs) from the Unified Human Gastrointestinal Protein catalogue.

Input: uhgp_50_mags.faa file | UHGP-50 MAG protein sequences
Databases: PDB100 → AFDB UniProt v4 → ESM highquality_clust30 | Three-database hierarchical search |
Skip PDB: No | PDB100 searched first
MMseqs2 parameters:

- Sensitivity | 5.7 (default)
- Max E-value: 0.001 (default)
- Min identity: 0.0 | No identity filtering at MMseqs2
- Min coverage: 0.9 (default)
- Top-K | 0 (all hits) | All passing hits retained

Post-filtering:

- Identity bins: [0.00, 1.01) | Single bin covering all identities
- Per-query selection: `topbits` (default) | Best hit per query by bitscore
- Drop self-hits: No | Self-hits kept (MAGs are unlikely to be in the database)

PyOpal alignment filtering:

- cmap-identity: 0.2 | Alignments with < 20% identity discarded
- Cmap-coverage: 0.9 | Alignments with < 90% query coverage discarded

Contact map:

- Generate contacts: 2

Prediction:

- Processing modes: BP, CC, MF | Biological Process, Cellular Component, Molecular Function
